# Identifying Human Encounters that Shape the Transmission of *Streptococcus Pneumoniae* and Other Respiratory Infections

**DOI:** 10.1101/116079

**Authors:** Olivier le Polain de Waroux, Stefan Flasche, Adam J Kucharski, Celine Langendorf, Donny Ndazima, Juliet Mwanga-Amumpaire, Rebecca F Grais, Sandra Cohuet, W John Edmunds

## Abstract

Although patterns of social contacts are believed to be an important determinant of infectious disease transmission, there is little empirical evidence to back this up. Indeed, no previous study has linked individuals’ risk of respiratory infection with their current pattern of social contacts. We explored whether the frequency of different types of social encounters were associated with current pneumococcal carriage and self-reported acute respiratory symptoms (ARS), though a survey in Uganda in 2014. In total 566 participants were asked about their daily social encounters and about symptoms of ARS in the last two weeks. A nasopharyngeal specimen was also taken from each participant. We found that the frequency of physical (i.e. skin-to-skin), long (≥1h) and household contacts – which capture some measure of *close* (i.e. relatively intimate) contact –, was higher among pneumococcal carriers than non-carriers, and among people with ARS compared to those without, irrespective of their age. With each additional physical encounter the age-adjusted risk of carriage and ARS increased by 6% (95%CI 2-9%) and 9% (1-18%) respectively. In contrast, the number of *casual* contacts (<5 minutes long) was not associated with either pneumococcal carriage or ARS. A detailed analysis by age of contacts showed that the number of close contacts with young children (<5 years) was particularly higher among older children and adult carriers than non-carriers, while the higher number of contacts among people with ARS was more homogeneous across contacts of all ages. Our findings provide key evidence that the frequency of *close* interpersonal contact is important for transmission of respiratory infections, but not that of *casual* contacts. Such results strengthen the evidence for public health measures based upon assumptions of what contacts are important for transmission, and are important to improve disease prevention and control efforts, as well as inform research on infectious disease dynamics.

**Author summary:** Although social contacts are an important determinant for the transmission of many infectious diseases it is not clear how the nature and frequency of contacts shape individual infection risk. We explored whether frequency, duration and type of social encounters were associated with someone’s risk of respiratory infection, using nasopharyngeal carriage (NP) of *Streptococcus pneumoniae* and acute respiratory symptoms as endpoints. To do so, we conducted a survey in South-West Uganda collecting information on people’s social encounters, respiratory symptoms, and pneumococcal carriage status. Our results show that both pneumococcal carriage and respiratory symptoms are independently associated with a higher number of social encounters, irrespective of a person’s age. More specifically, our findings strongly suggest that the frequency of *close* contacts is important for transmission of respiratory infections, particularly pneumococcal carriage. In contrast, our study showed no association with the frequency of short *casual* contacts. Those results are essential for both improving disease prevention and control efforts as well as informing research on infectious disease dynamics and transmission models.

## Introduction

The transmission of respiratory infections is likely to depend on the frequency and age structure of human social contacts, as well as other factors including pre-existing immunity from prior infection or vaccination [1-3]. To understand the dynamics of such infections, studies to quantify social mixing patterns have been conducted in various settings, under the assumption that self-reported encounters reflect transmission probabilities of pathogens transmitted through close contact [2, 4-11]. Combined with disease transmission models, these data are increasingly being used to inform infection control policies [12, 13].

There is evidence from population-based models that self-reported social mixing patterns can reproduce observed aggregated seroprevalence data for chickenpox [14], mumps [4], parvovirus [15], influenza [4, 16, 17]) and whooping cough [18]. Moreover, it has been suggested that age-stratified social mixing patterns can capture individual influenza risk, as measured by a four-fold rise in neutralization titres over the course of an epidemic (17, 18). However, it remains unclear precisely how risk of infection is related to the frequency and nature of an individual’s social encounters around the time of infection.

To establish how social behaviour shapes individual-level infection, we explored whether the frequency and duration of different types of social encounters were associated with an individual’s risk of respiratory infection, using nasopharyngeal (NP) carriage of *Streptococcus pneumoniae* (the pneumococcus) and self-reported acute respiratory symptoms (ARS) as endpoints. *S.pneumoniae* is one of the main causes of pneumonia and sepsis globally [19], disproportionally so in low-income settings [19-21]. Colonization of the nasopharynx is a precondition to disease, and the main source of human-to-human transmission. Given that most episodes of carriage remain asymptomatic, social behaviour is unlikely to change as a result of carriage, making pneumococcal carriage a more suitable endpoint than symptomatic illness to explore the association between social behaviour and infection risk, given that people tend to limit their contacts during symptomatic illness [22]. In addition, as natural immunity to carriage is weak [23], pre-existing immunity is less likely to confound associations between disease and social contact patterns than in studies using immunizing infections as biological endpoint [16, 17].

We analysed data from a social contact study nested within a survey of NP carriage of *S.pneumoniae* conducted across a rural South-West Uganda in 2014, before the introduction of the pneumococcal conjugate vaccine (PCV). Participants across age groups were randomly selected from 56 villages and one small town. Participants were asked to name all individuals with whom they had a conversational encounter lasting ≥ 5 minutes between wake-up on the day prior to the survey day to wake-up on the survey day (∼ 24 hours). Those encounters are further referred to as *‘ordinary’* contacts. For very short contacts (i.e. <5 minutes), which we here define as *‘casual’* contacts, an estimate of the number of contacts was asked (<10, 10-19, 20-29, ≥30) without further details. ARS were defined as any of the following symptoms in the previous two weeks: sore throat, sneezing, difficulty breathing, and runny nose. A nasopharyngeal specimen was taken from each survey participant and was processed and analysed as per WHO guidelines [24]. Using these individually-matched data, we examined whether the type and frequency of social contacts differed by carriage status and between individuals with and without ARS.

## Results

### Study population

Of the 687 individuals initially targeted for inclusion, 568 (83%) individuals from unique households responded to the survey, with data on both social contacts and carriage or ARS available from 566 participants. Participant’s age spanned across age groups and the sex distribution was reasonably balanced (58% female). The majority (98%) of children aged 6 – 15 years attended school or college. More than a third of all adults (36%) worked in agriculture and 22% were homemakers/housewife.

On average, people made seven *‘ordinary’* contacts (defined as contacts ≥5 minutes long), ranging from 0 to 25, the majority of which were physical (i.e. ‘skin-to-skin’ contact or indirect physical contact through utensils passed from mouth-to-mouth). There was no evidence that the average number of contacts differed by weekday or between weekdays and weekends (P=0.623). Children aged 5-9 years reported most contacts and children <5 years the fewest (Figure 1). The majority of contacts made by children were physical.

**Figure 1:**
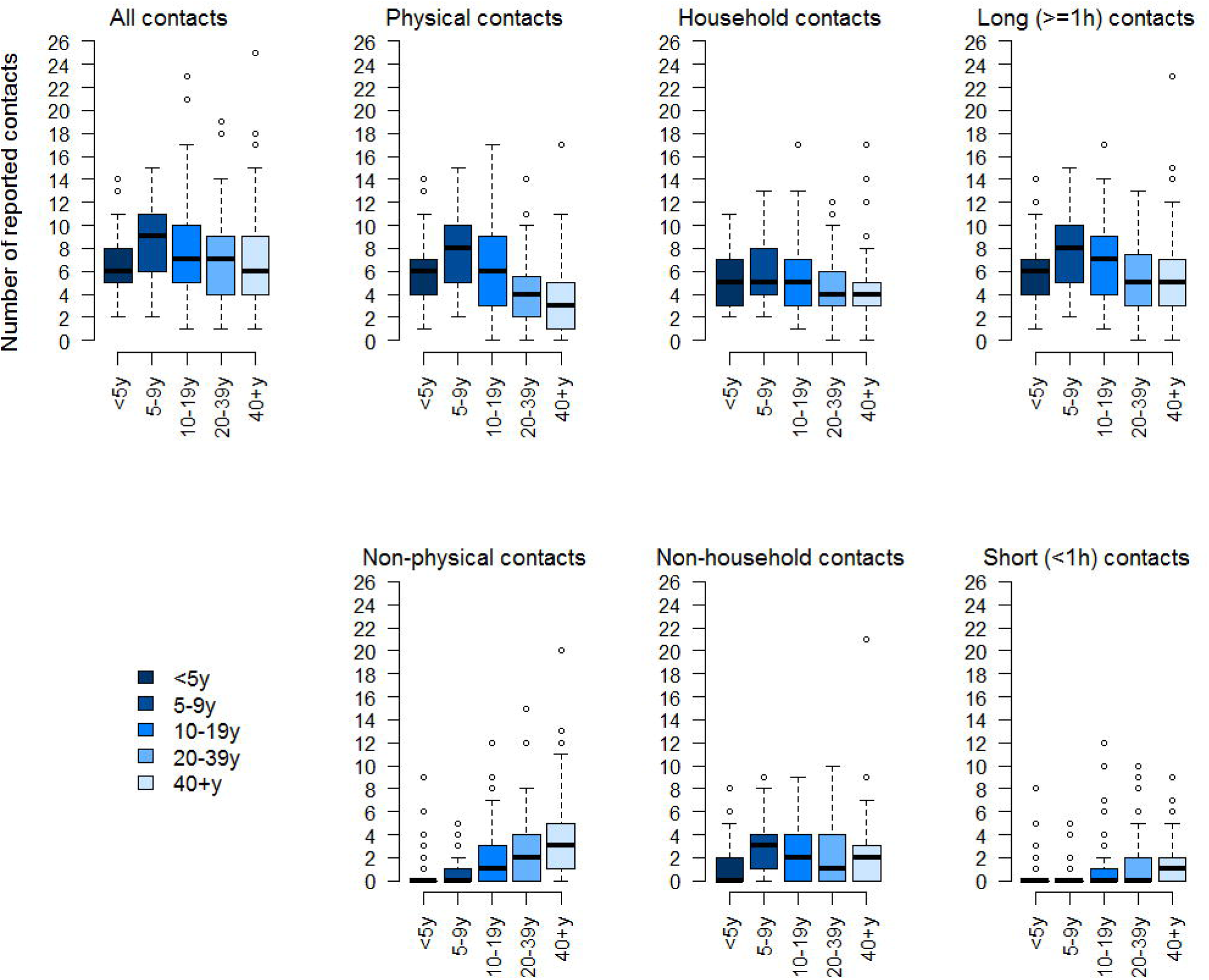
Number of reported contacts by age group and type of contact. <u>Legend</u>: Boxplots showing the median (horizontal line), interquartile range (boxes) and 95% confidence interval around the mean (whiskers). The outliers are shown as bubbles above or under the whiskers

The most intense mixing tended to be between individuals of the same age group (i.e. assortative mixing), but there was also substantial mixing between age groups. Contact from and with children aged <10 years involved proportionally more physical touch than contacts between older children and adults (Figure 2A and 2B). The proportion of non-physical contacts, contacts outside the household and contacts of short (<1h) duration was higher among teenagers and adults than among younger children and infants.

**Figure 2:**
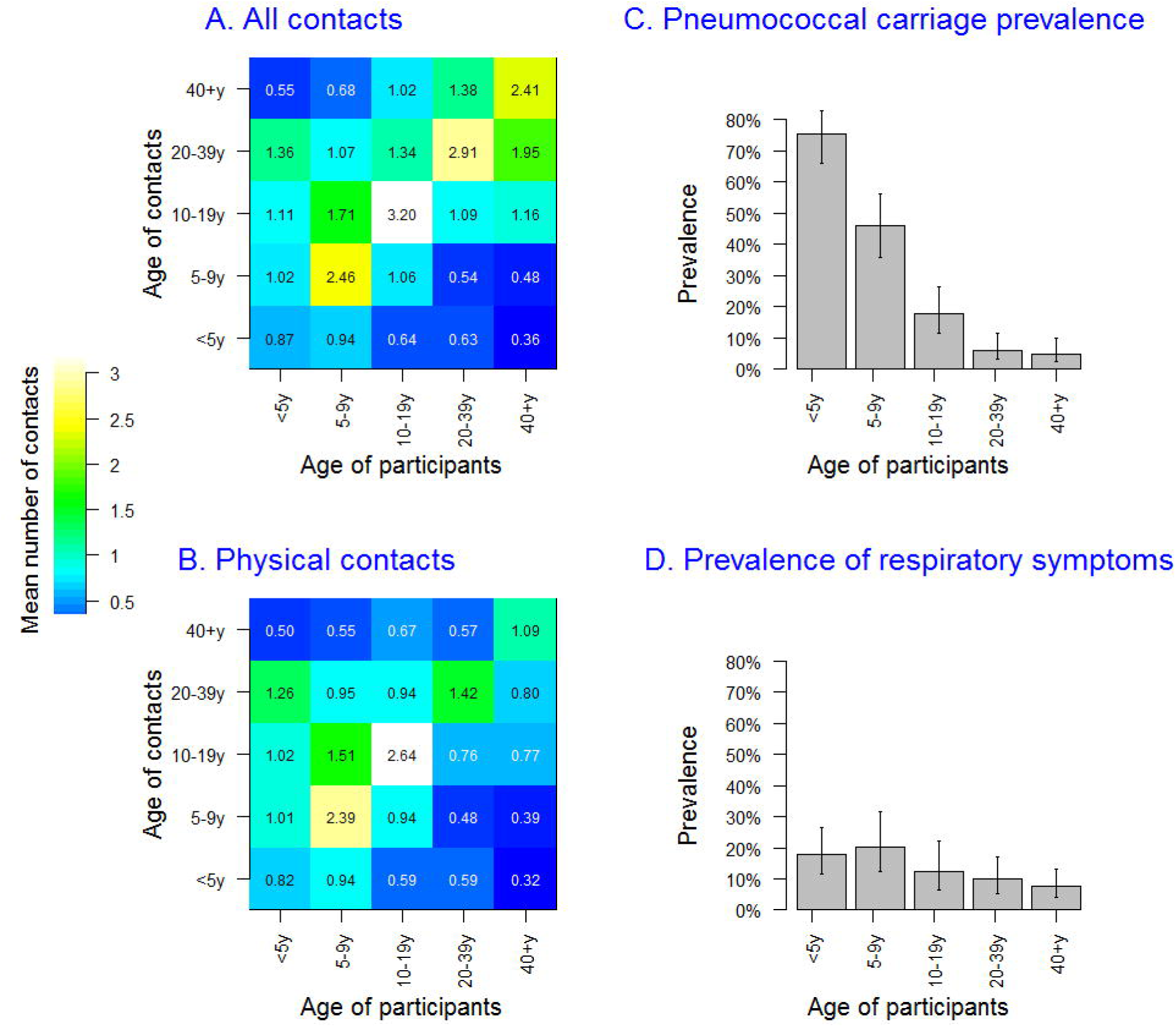
Contact matrices and prevalence of *S.pneumoniae* carriage and Acute Respiratory Symptoms (ARS), by age. Legend: Contact matrices and prevalence of *S.pneumoniae* carriage (Panel A) and Acute Respiratory Symptoms (Panel B). In panels C and D, the height of each bar corresponds to the point prevalence and the error bar represents the 95% Confidence Interval (CI).

There was no difference in contact patterns by sex, for all types of contacts considered.

Four hundred and ninety (87%) participants estimated how many casual contacts (i.e. <5 minutes in duration) they had the day before the survey. Over a third (36%) of these reported 10 or more casual contacts, and 11% reported more than 20 casual contacts. Among the 13% who could not estimate how many casual contacts they had, there were proportionally more children with over half (56%) under 10 years. The mean number of reported ‘ordinary’ contacts was higher among individuals reporting ≥10 casual contacts than those reporting fewer than 10 contacts (mean 8.9 vs 5.9, P<0.001), as well as among the 78 individuals who did not know how many casual contacts they had (mean 8.8, P<0.001).

The prevalence of pneumococcal carriage was strongly age dependent, decreasing from 75% in children <5 years to 46% among 5 – 9 year olds, 17% in 10 – 19 year olds, and further decreasing to 8% and 7% among 20-39 years and ≥40 years old respectively (Figure 2C). Overall, 72 (13%) people reported having suffered from ARS in the two weeks prior to the survey. The prevalence of ARS varied much less with age than that of carriage (Figure 2D), ranging from 20% among 5 to 9 year olds to 8% among ≥40 years old. There was no sex difference in the prevalence of carriage (age-adjusted RR for males 0.97, P=0.814) or ARS (age-adjusted RR for males 1.40, P=0.134). Carriage and ARS were poorly correlated (Pearson’s correlation coefficient R=0.09, ranging from -0.08 to 0.29 by age group), and there was no evidence that the risk of ARS was higher among carriers compared to non-carriers (age-adjusted relative risk (RR) 1.06 (95%CI 0.85 – 1.33)).

### Social contacts as a risk for pneumococcal carriage or ARS

Overall, the mean number of contacts among carriers was significantly higher than non-carriers, and this observation was consistent across age groups, although most differences were not statistically significant due to small numbers (Figure 3). In particular, carriers had more physical contacts (Fig 3A). This pattern was also consistent for ARS, with the exception of 5 – 9 year olds in whom the mean number of contacts among individuals with ARS was lower for physical contacts (Fig 3B).

**Figure 3:**
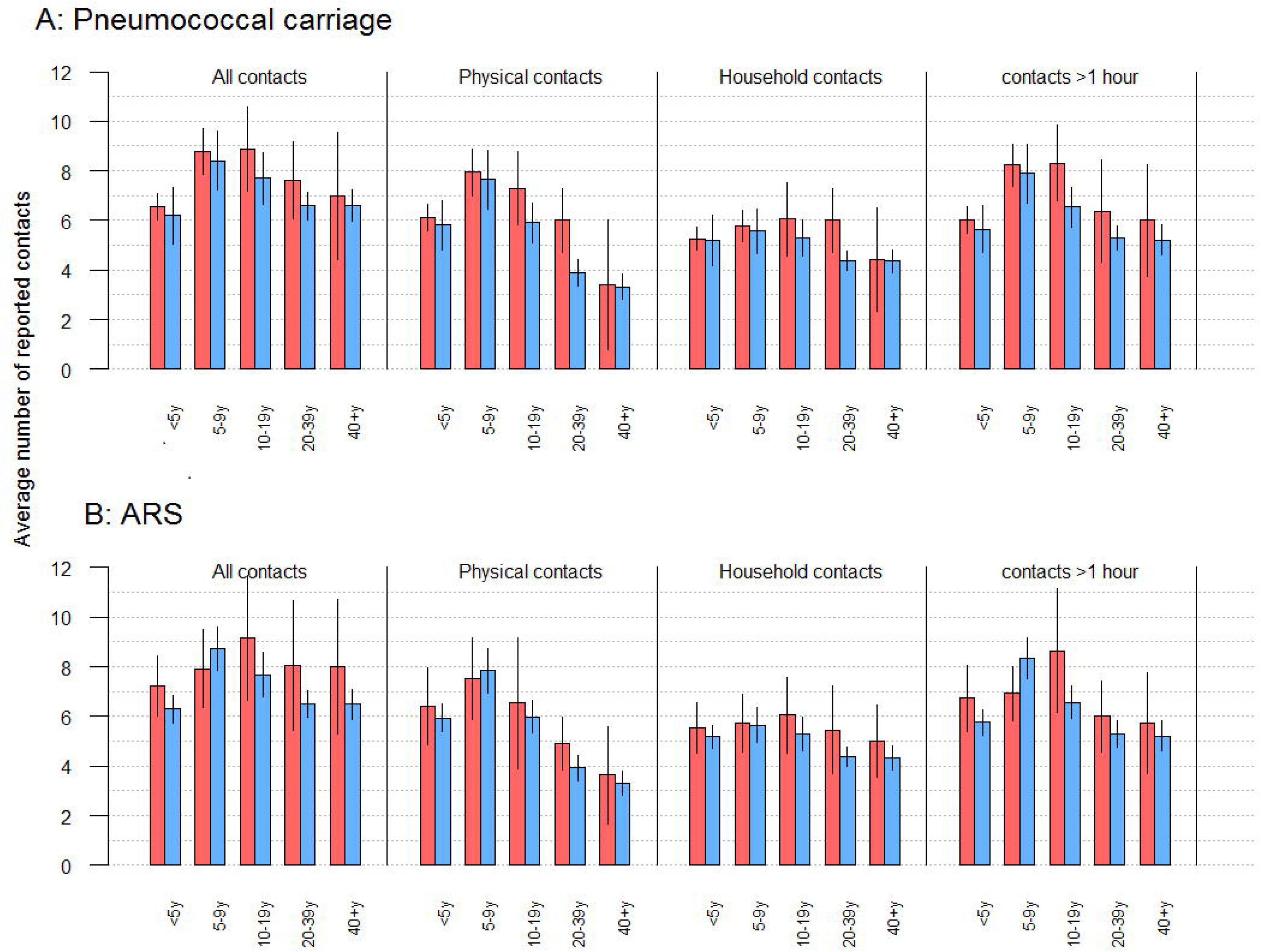
Mean number of contacts by age group and nasopharyngeal carriage status (Panel A) or ARS status (Panel B). Legend: The graph shows the mean number of contacts among carriers (red, panel A), non-carriers (blue, panel A), individuals reporting ARS (red, panel B), and individuals without ARS (blue, panel B). This is shown by age group and for all contacts, physical contacts, household contacts and contacts lasting over 1 hour.

In univariable analysis, the risk of carriage increased with all contacts, household contacts, contacts ≥1 hour long and physical contacts. The latter had the largest effect size, with a 13% increased risk for each additional contact reported by participants (Table 1). Physical contacts and contacts ≥1 hour were strongly correlated (R = 0.76), particularly among children <5 years (R=0.85) and children aged 5 – 9 years (R=0.87), hence their effect could not be disentangled, whereas the correlation between physical contacts and household contacts was moderate (R=0.61, ranging from 0.47 to 0.70 between age groups). After age adjustment, physical contacts or contacts ≥1 hour remained most significantly associated with carriage, with a 6% increased risk (95%CI 2 – 10%) for each unit increase in the number of reported contacts (Table 1). We found good evidence that the number of household contacts increased the risk as well (Table 1). There was no confounding effect by other covariates and models were therefore only adjusted for age (Supporting Information S1).

An increase in physical, household and long (≥ 1h) contacts, were also associated with an increased risk of ARS in univariable analysis. There was little or no evidence of a confounding effect of age, given the more constant prevalence of ARS across age groups (Table 1). Unlike pneumococcal carriage, however, the relative risk was more constant across types of contacts, and household contacts rather than physical contacts were most strongly associated with a risk for ARS, with a risk increase of 9% (1 – 18%) for each additional reported contact.

**Table 1:** Relative risk of ***S.pneumoniae* carriage and ARS** by frequency of contact

| Contact types | Mean contacts |  | Crude RR (95%CI) | Adjusted RR (95%CI) |
| --- | --- | --- | --- | --- |
| NP carriage | NP+ | NP- |  |  |
| All contacts | 8.09 | 7.53 | 1.04 (1.00; 1.07)* | 1.03 (1.00; 1.07)* |
| Physical contacts | 6.83 | 6.09 | 1.13 (1.09; 1.17)* | 1.06 (1.02; 1.09)* |
| Non-physical contacts | 1.25 | 1.43 | 0.78 (0.72; 0.86)* | 0.97 (0.91; 1.05) |
| Household contacts | 5.64 | 5.31 | 1.09 (1.03; 1.14)* | 1.04 (0.99 ; 1.10) |
| Non-household | 2.22 | 2.45 | 0.98 (0.93; 1.04) | 1.03 (0.98; 1.09) |
| Contacts ≥ 1hour | 7.50 | 6.56 | 1.08 (1.05; 1.12)* | 1.06 (1.02; 1.10)* |
| Contacts <1h | 2.38 | 2.62 | 0.81 (0.73; 0.92)* | 0.94 (0.85; 1.03) |
| ARS | ARS+ | ARS- |  |  |
| All contacts | 8.18 | 7.63 | 1.07 (1.02; 1.12)* | 1.07 (1.02; 1.13)* |
| Physical contacts | 6.10 | 6.26 | 1.09 (1.03; 1.15)* | 1.07 (1.00; 1.14)* |
| Non-physical contacts | 2.07 | 1.36 | 1.00 (0.89; 1.10) | 1.06 (0.95; 1.18) |
| Household contacts | 5.70 | 5.31 | 1.10 (1.02; 1.19)* | 1.09 (1.01; 1.18)* |
| Non-household | 2.47 | 2.31 | 1.04 (0.98; 1.11) | 1.05 (0.98; 1.13) |
| Contacts ≥ 1hour | 7.33 | 6.74 | 1.07 (1.01; 1.13)* | 1.07 (1.00; 1.14)* |
| Contacts <1 hour | 3.14 | 2.58 | 1.05 (0.96; 1.15) | 1.08 0.99; 1.18) |

Legend: \*=associations significant at P<0.05. RR= Risk Ratio. CI= Confidence Interval. NP+= pneumococcal carrier, NP - = non-carrier, ARS+ = with ARS, ARS- = without ARS

Next, we analysed whether the number of casual contacts (i.e. contacts lasting < 5 minutes) was associated with either pneumococcal carriage or ARS. We found no evidence that the prevalence of pneumococcal carriage or the risk of ARS were associated with reporting higher levels of casual contacts, as shown in Table 2. Given the small numbers of individuals reporting ≥30 casual contacts, we pooled the 56 individuals reporting ≥20 casual contacts into one category.

In univariable analysis the risk of pneumococcal carriage was higher in the group of 78 participants who did not know how many casual contacts they may have had. This was due to the higher proportion of children <5 years and aged 5 – 9 years in that group, however, after age-adjustment, there was no evidence that the risk of pneumococcal carriage was higher in that group. Similarly, a higher number of casual contacts was associated with increased risk of ARS, and with no confounding effect of age or other variables on the estimates.

We found no confounding effect on casual contacts on the relative risk of carriage or ARS as a function of the frequency of reported ‘ordinary’ contacts.

**Table 2:**
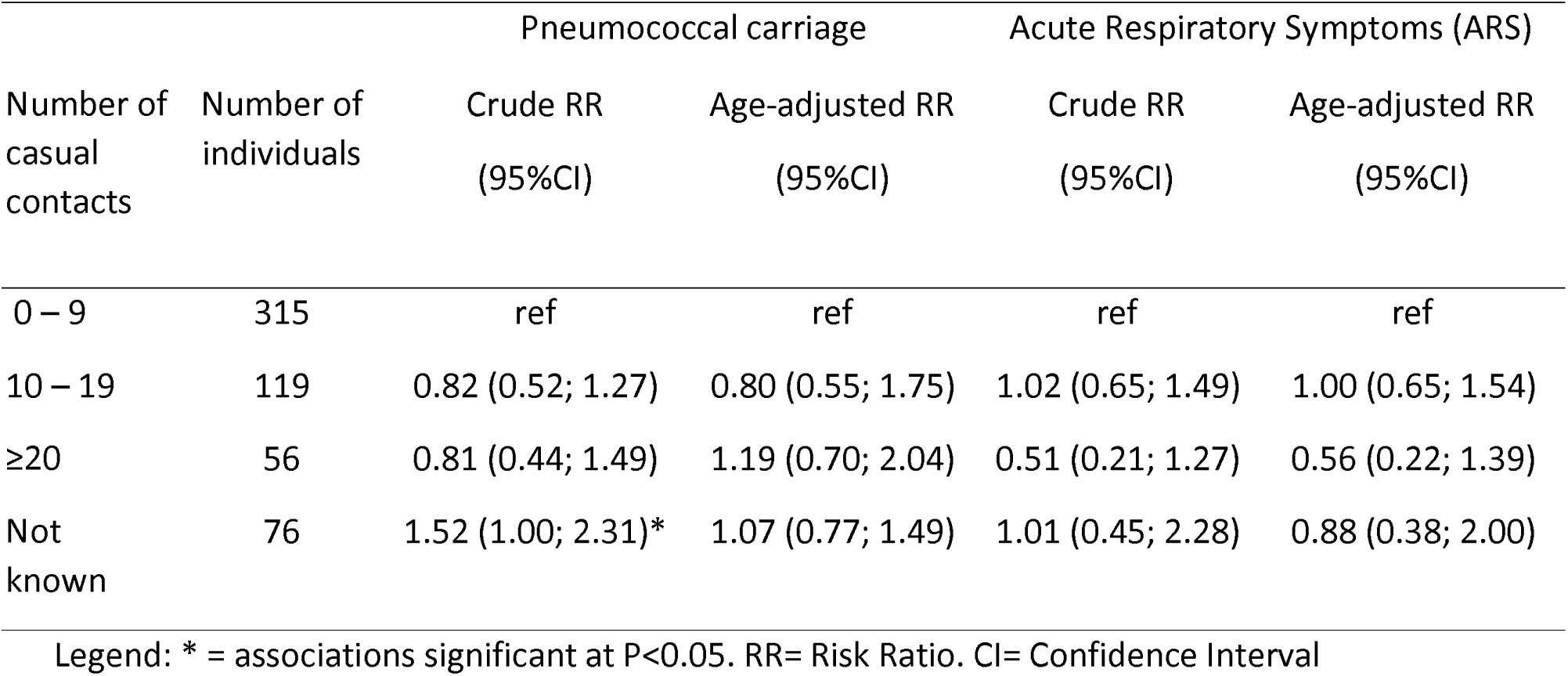
Relative risk (RR) of pneumococcal carriage and ARS by level of reported number of daily casual contacts

Finally, we explored the characteristics of age-specific mixing patterns by ARS or carriage status in greater detail, with a focus on physical contacts and household contacts. We computed the ratio of the mean number of contacts within and between age groups among carriers compared to non-carriers, and among individuals with ARS compared to those without. The average number of physical encounters within and between age groups tended to be higher for carriers than non-carriers in most instances, albeit with substantial uncertainty owing to small numbers (Figure 4). Carriers reported more contacts with children < 5 years, and particularly adult carriers who reported on average more than twice as many physical contacts with children under five than non-carriers (Figure 4). Given that most of such contacts occurred within the household, the effect of physical and household contacts with children <5 years was indistinguishable, whereas for older age groups, the association with physical contact was stronger than that with household contacts (Figure 4). Similar findings were seen for ARS, however, unlike for pneumococcal carriage, symptomatic adults did not have more contacts with young children than asymptomatic ones. Results based on absolute differences in the mean number of contacts rather than relative means are displayed in Figure S1 (Supporting Information), showing the similar associations than the reported ratios, but providing a quantified difference in mean number of contacts instead.

**Figure 4:**
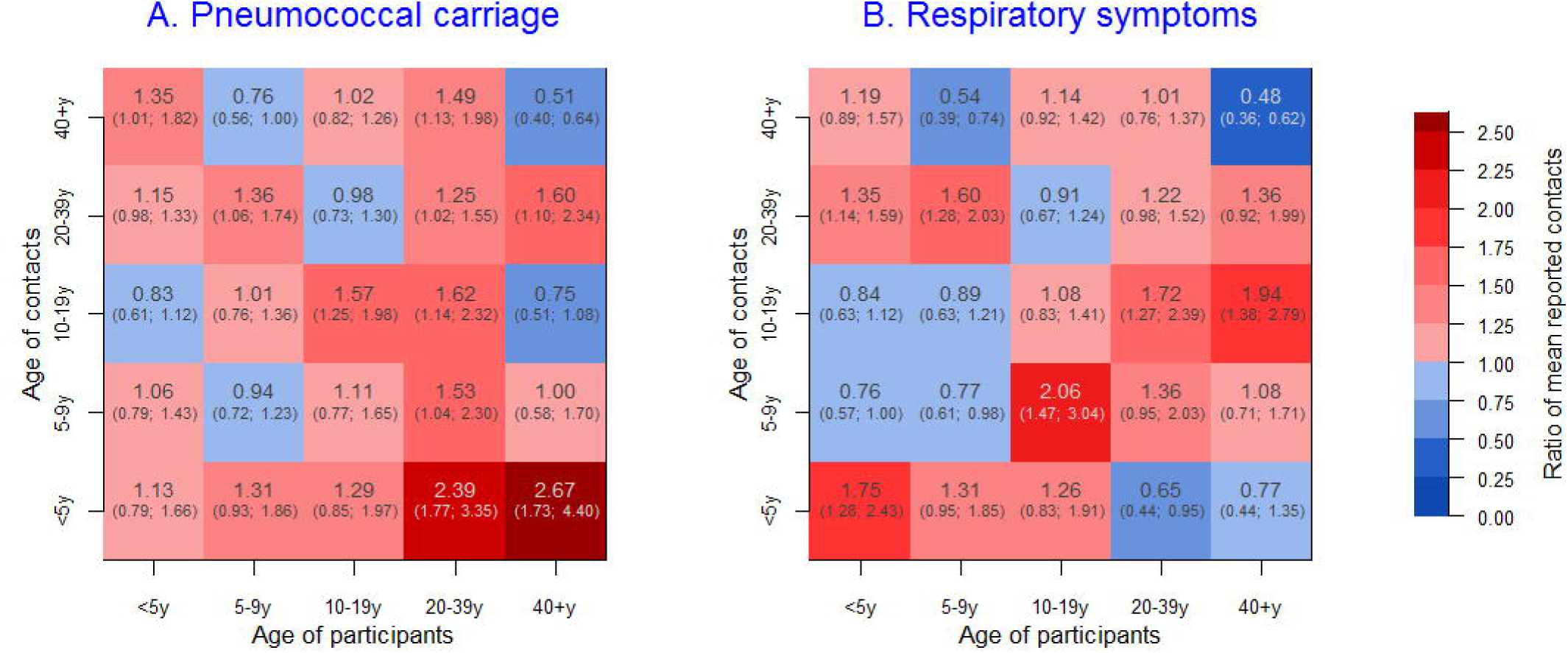
Ratio of the mean reported number of *physical* contacts between pneumococcal carriers and non-carries (panel A) and individuals with ARS compared to those without ARS (Panel B). Legend: Matrices of the ratio of the mean number of contacts among pneumococcal carriers compared to non-carriers (Panel A) and individuals with ARS compared to non-ARS (panel B). The numbers represent the point estimate of the ratio and in brackets the 95% confidence bounds. Blue are cells with a point estimate <1 and in red with point estimates >1.

## Discussion

Our study provided a unique opportunity to explore whether and how social contact patterns are associated with someone’s risk of acute respiratory infection in a mostly rural East African setting. Our results show that people who tend to have more frequent close contacts are more likely to be pneumococcal carriers or to report acute respiratory symptoms, irrespective of their age. In contrast, we found that less intimate or short casual contacts were not associated with someone’s infection risk, suggesting that social contacts important for transmission are close interpersonal encounters.

Existing evidence on the association between contact patterns and risk of infection is mostly ‘ecological’, with very few studies based on individual-level data. Using data of influenza A/H1N1 seroconversion in Hong Kong coupled to social contact data in the same population, Kwok et al. [16] showed that age rather than social contact patterns were the main driver of the individual risk of infection in that setting, and further work by Kucharski et al. [17] on the same data further supported the finding that someone’s risk of infection is related to the average mixing pattern within their age group rather than their reported number of contacts. However, the validation of the ‘social contact hypothesis’ with such data remains difficult, due to challenges in accounting for acquired immunity in the population, assumptions around stability of behaviour over relatively long time periods as well as challenges in capturing influenza infection events based on serological data only [16, 17].

Studying nasopharyngeal carriage *of S.pneumoniae* as the main biological endpoint enabled us to address many of these issues, for several reasons. First, the high prevalence of carriage provided the statistical power required to study individually-matched acquisition risk in our study. Second, a person’s individual behaviour is unlikely to be affected by carriage given that the vast majority of episodes remain asymptomatic, in contrast to symptomatic respiratory infections during which social behaviour might change as a result of illness [22]. Most carriage episodes in our study were asymptomatic, and our decision to assess the risk for ARS and NP carriage separately stemmed from that observation. In addition, given that duration of carriage is relatively short – at most 3 months in young children and no longer than a few weeks in adults [25] –, and as no difference in our survey was observed in the mean number of contacts between days of the weeks and between survey weeks, it is reasonable to assume that contact patterns measured on a given day reflect contact patterns around the time of pneumococcal acquisition. Finally, as colonisation results in weak protective immune responses and limited reduction in serotype-specific reacquisition risk [23, 26], and with over 90 circulating serotypes, natural acquired immunity is unlikely to distort the association between carriage and social contact patterns, unlike immunizing infection for which individuals with more frequent risk of infection due to their social contact patterns are also more likely to be immune.

In contrast, self-reported respiratory symptoms may be influenced by factors such as behaviour change in illness [22] or immunity. The definition itself also presents limitations; although respiratory viruses such as the Respiratory Syncytial Virus (RSV), adenovirus, parainfluenza and influenza viruses are likely to account for a large proportion of ARS cases [27], the definition was used as a non-specific proxy for acute respiratory infection and may have captured other infectious and non-infectious conditions. Notwithstanding such caveats, results for ARS were similar to those for nasopharyngeal carriage, but also showed a more consistent association with all types of non-casual contacts, rather than physical or contacts of long duration only. This suggests that the definition may have mostly captured acute infections, and also provides further evidence that the number of close interpersonal contacts, and particularly household contacts, plays a role in the transmission of acute respiratory pathogens.

One of the striking features of our analysis of the relative number of mean contacts between age groups is that adults colonised with *S.pneumoniae* reported more than twice as many close contacts with children under five than non-carriers. This is in agreement with observational and modelling studies showing that pneumococcal carriage risk increases with household size and with the number of children <5 years in the household [28]. It also supports the finding that carriage acquisition in adults occurs mostly within the household as a result of contact with young children who are drivers of infection [29]. In contrast, however, we found that adults with ARS reported an equal or lower number of contacts with children <5 years than asymptomatic adults. This likely reflects the epidemiological differences between carriage and ARS, given the very high prevalence of carriage among <5 year olds and the much less marked difference in age-specific prevalence of ARS across age groups, in addition to other potential factors such as acquired immunity among adults more frequently exposed to very young children. Some of the specific details of the contact patterns and differences between *S.pneumoniae* and ARS are harder to interpret, due in part to wide statistical uncertainty, particularly among adults in whom the number of carriers or symptomatic individuals is small. However, overall the results from our analysis support the general finding that increased close contacts are associated with higher risk of ARS or pneumococcal carriage.

There is still much debate about how respiratory pathogens are transmitted from person to person; whether through close direct or indirect physical contact, through large droplet transmission at close range (<1 meter), or through aerosolized particles floating over longer distances, particularly in poorly ventilated indoor settings [30, 31]. It is likely that many pathogens can be transmitted through a combination of these routes. Yet the contribution of each mechanism remains uncertain [30, 31]. It is generally assumed that the main transmission route of *S.pneumoniae* is through direct contact [32], as well as indirectly through shared glasses or bottles [33]. Analogously, for other colonising bacteria such as *N.meningitidis*, close contact and intimate kissing are known risk factors among teenagers and young adults [34]. Similarly, it is believed that influenza and other respiratory viruses are primarily transmitted through direct contact or contact with large droplet transmission at close range rather than aerosolized particles [30].

Although our objective was not to demonstrate transmission, our findings strongly support that direct close interpersonal contact is an important mode of transmission for pathogenic bacteria of the nasopharynx as well as respiratory viruses, and strengthens the scientific evidence for public health measures based upon these assumptions, such as hand washing campaigns or chemoprophylaxis of close contacts of cases of meningococcal meningitis to name a few. The strong association of ARS with household and other close contacts, independently from physical touch, might suggest that other mechanisms such as indirect transmission through fomites or aerosol transmission may play a role. Elucidating the contribution of such factors in this context would be an important question for future research.

Our findings have also several implications for infectious disease research. Contact structures are central to transmission models, and appropriate assumptions about what type of contact drives infectious disease transmission are essential. Our results suggest that the parameterisation of transmission models of *S.pneumoniae* and similar pathogens using mixing matrices based on physical or another measure of *close* interpersonal contact would more likely capture relevant contact patterns than those based on any type of social encounter. This has also implications for the design of contact studies, particularly in low-income settings given the scarcity of published data currently available and the need to collect additional data from many more settings [7, 8]. In contrast to diary-based approaches that have been used by many [2, 4, 7, 8], the study design here was relatively simple and only involved a single face-to-face interview. A drawback of such retrospective approach is the lack of detailed information about very short (i.e. ‘casual’) contacts, as such information was deemed unreliable – and this is further supported by evidence that short contacts tend to be inconsistently recorded even in prospective diary-based approached [1]. However, given that very short contacts may not account for much of the transmission, as our study suggests, a more simple retrospective design is a potential attractive option in other settings where contact data are lacking, and in which data collection through more comprehensive diary-based approaches may be difficult to implement.

There are some additional limitations to our study. We were unable to explore other potential confounding factors, such as bed share, ventilation, indoor smoke or hand washing [35], and the contribution of such factors should be explored further. Moreover, it remains unclear to what extent our results can be generalised to any acute respiratory infection. For example, factors such as ventilation and airflow may be of particular importance for aerosolized transmission of pathogens such as mycobacteria [7], compared to influenza or *S.pneumoniae* [30, 31], and whether household or physical contacts reflect contact patterns important for aerosolized transmission remains uncertain [7]. Finally, while we found a strong association at the individual level, our study does not demonstrate causality.

However, in the absence of detailed longitudinal data on acquisition events between all individuals’ contacts —which would be challenging and possibly unrealistic to obtain— our findings provide consistent evidence of a ‘dose-response’ association at the individual level between close social encounters and acquisition risk for respiratory pathogens, and therefore provides robust support both for the social contact hypothesis, and for research and policy work based upon this hypothesis.

## Materials and Methods

### Data collection

The study was conducted in Sheema North Sub-District (Sheema district, South-West Uganda) between January and March 2014. Sixty clusters were randomly selected from the 215 villages and two small district towns (Kabwohe and Itendero) in the study area, proportionally to the population size of each village and town. In each cluster 11 or 12 individuals were randomly sampled from different households to both answer questions about their social contacts and their history of respiratory illness in the last two weeks, as well as having a nasopharyngeal swab taken. A household was defined as the group of individuals living under the same roof and sharing the same kitchen on a daily basis.

For the social contact questionnaire, participants were first asked to list all the individuals with whom they had a two-way conversational contact lasting for ≥5 minutes during a period of approximately 24 hours prior to the survey day (from wake up the previous day until wake up on the survey day). Such encounters were defined as ‘ordinary contacts’. For each reported ordinary contact, participants (or their parent/guardian) were asked to estimate the contact’s age (or estimated age), how long the encounter lasted for and whether it involved skin-to-skin touch or utensils passed from mouth to mouth (either of those defining ‘physical contacts’). For very short social encounters (<5 minutes), which were defined as ‘casual contacts’ (e.g. seeing someone on the way, encounter in a shop etc.), participants were asked to estimate the number of encounters based on pre-defined categories (<10 contacts, 10-19 contacts, 20-29 contacts, ≥30 contacts), but not to provide further details about each contact.

Next, participants were asked about respiratory symptoms experienced in the two weeks prior to the survey, including any of the following: cough, runny nose, sneezing, sore throat, difficulty breathing.

Finally, after the interview was completed, a nasopharyngeal swab was taken from each participant. NP samples were collected, transported and analysed as per WHO guidelines [24]. NP swabs (flocked nylon swabs, COPAN, Italy) were inoculated in a skim milk tryptone-glucose-glycerol (STGG) medium, transported in cool boxes and frozen at the research laboratory at -20°C within 8 hours of collection. Specimens were inoculated onto a selective agar plate of 5 mg/L gentamicin-Columbia agar with 5% sheep blood and incubated at 37°C in 5% CO_2_ atmosphere overnight. Pneumococcal identification was based on optochin susceptibility testing of all alpha-hemolytic colonies and bile solubility testing in case of intermediate susceptibility to optochin.

### Statistical analysis

We first performed descriptive analyses, with age-specific probability weights to account for different inclusion probabilities by age at the design stage, and adjusted for the clustering by village through the use of clustered ‘sandwich’ variance estimators to account for possible correlation within each of the sixty clusters [36]. We explored social contact patterns within and between age groups through contact matrices. In such matrices we report the mean number of ordinary contacts of participants in age group *j* with contacts in age group *i* (*m_ij_*), adjusted for reciprocity, as in Melegaro et al. [15].

We then analysed whether and how the frequency distribution of contacts was associated with pneumococcal carriage or self-reported ARS.

We modelled the effect of ‘ordinary’ contacts (defined as contacts ≥5 minutes long) on carriage or respiratory symptoms as a function of contact frequency, through a Poisson model with a robust variance estimator, and inclusion probability weights by age group. We treated contacts as continuous variables, but assessed departure from linearity through likely ratio tests and model comparisons of Bayesian Information Criterion (BIC). In multivariable analysis, we considered for inclusion any covariate significantly associated with the outcome at P<0.10 in univariable analysis. Model improvement was considered for any decrease in the BIC. Further details are provided in the Supporting Information S1.

Next, we used the same analytical approach to assess whether carriage and ARS were associated with the level of self-reported ‘casual’ contacts (i.e. <5 minutes long).

Finally, we explored differences in the social mixing matrices by status of pneumococcal carriage and ARS. To do so, we computed the ratio of the mean number of reported contacts by participants *j* with contacts *i* among carriers compared to non-carriers 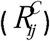 or symptomatic compared to asymptomatic individuals 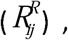 such that 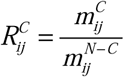 and 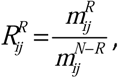, where *C* = carriers, *R* = participants with ARS and *N* = the total number of participants. Estimates were not adjusted for reciprocity given that subpopulations were not closed (e.g. contacts of carriers may not be carriers), and that our aim was to compare reported number of contacts rather than calculate a contact matrix. The uncertainty in reported values and ratios was obtained through resampling techniques, drawing random samples from each *m_ij_*, with the number of draws equal to the study population in each age group *j*. Ratios and uncertainty around them was obtained from the ratio of bootstrapped matrices.

The same approach as described above was used to compute the differences in *m_ij_* with 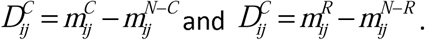

All analyses were performed in STATA 13.1 IC and R version 3.2.

### Ethics

Approval was obtained from the Ethical review boards of the London School of Hygiene and Tropical Medicine, Médecins Sans Frontières, the Faculty of Medicine Research & Ethics Committee of the Mbarara University of Science and Technology, the Institutional Ethical Review Board of the MUST, and the Uganda National Council for Science and Technology.

## Acknowledgments

We thank all the surveyors for their work and all the individuals who took part in this study, Rinah Arinaitwe for her help in managing the data entry, Dan Nyehangane for his support with the laboratory analyses, and Dr Yap Boum II for his help overseeing field activities. We are also grateful to Prof Anthony Scott, Dr Fabienne Nackers and Prof Francesco Checchi for their valuable input in the design and protocol of the field surveys.

## Supporting Information Captions

**SI Text.** Analytical approach and model comparison

**S1 Figure:** Difference in the mean number of *physical* contact between pneumococcal carriers and non-carriers (Panel A) and between individuals suffering and not suffering from respiratory symptoms (Panel B).<u>Legend</u>: Matrices of the mean difference between the mean number of contacts among pneumococcal carriers and non-carriers (panel A), and individuals with ARS compared to non-ARS (panel B). The numbers represent the point estimate and the lower and upper bound of the 95% credible interval are shown inside the brackets.

**S2Text:** Data dictionary **S3 Data:** Dataset

